# Post-translational covalent assembly of CAR and synNotch receptors for programmable antigen targeting

**DOI:** 10.1101/2020.01.17.909895

**Authors:** Jason Lohmueller, Adam A. Butchy, Yaniv Tivon, Michael Kvorjak, Natasa Miskov-Zivanov, Alexander Deiters, Olivera J. Finn

## Abstract

Chimeric antigen receptors (CARs) and synthetic Notch (synNotch) receptors are engineered cell-surface receptors that sense a target antigen and respond by activating T cell receptor signaling or a customized gene program, respectively. To expand the targeting capabilities of these receptors, we have developed switchable adaptor receptor systems for which receptor specificity can be directed post-translationally via covalent attachment of a co-administered antibody. Instead of directly targeting an antigen, our receptors contain the SNAPtag self-labeling enzyme, which reacts with benzylguanine (BG)-conjugated antibodies to assemble covalently-associated antigen receptors. We demonstrate that activation of SNAP-CAR and SNAP-synNotch receptors and their downstream effector functions can be successfully targeted by several clinically-relevant BG-conjugated antibodies in an antigen-specific and antibody dose-dependent manner. To better define parameters affecting receptor signaling, we developed a mathematical model of switchable receptor systems. SNAP receptors provide a powerful new strategy to post-translationally reprogram the targeting specificity of engineered cells.

## Introduction

Engineered antigen receptors are revolutionizing the treatment of blood cancers and show promise in cell therapies treating a wide range of other diseases^1^. The most clinically advanced of these technologies are chimeric antigen receptors (CARs), synthetic T cell receptors most often comprised of an antigen-specific antibody single chain variable fragment (scFv) fused by spacer and transmembrane domains to intracellular T cell signaling domains^2–4^. Upon binding to a target antigen, CARs stimulate T cell activation and effector functions including cytokine production, cell proliferation, and target cell lysis. Adoptively transferred CAR T cells targeting the B cell antigen CD19 are now FDA-approved and have been highly successful in treating refractory acute lymphoblastic leukemia^5–7^. Creating CARs against additional targets to treat other types of cancer and immune-related diseases is a major research focus^8,9^. Another class of highly versatile antigen receptors are synthetic Notch, “synNotch” receptors which consist of an antigen binding domain, the Notch core protein from the Notch/Delta signaling pathway, and a transcription factor ^10–12^. Instead of activating T cell signaling upon binding to the target antigen, the Notch core protein is cleaved by endogenous cell proteases thus releasing the transcription factor from the cell membrane. Subsequent nuclear translocalization leads to transcriptional regulation of one or more target genes. These receptors are highly modular, they can be created to target different cell surface antigens by changing the scFv, and can positively or negatively regulate any gene of interest by either fusing different transcription factors as components of the receptors or by changing the transgenes under their control. This versatile receptor type receives increasing clinical interest in immunotherapies as well as applications to tissue engineering^13,14^.

To gain additional control over CAR function, we and others have developed “switchable” adaptor CAR systems for which the CAR, instead of directly binding to an antigen on a target cell, binds to common tag molecule fused or conjugated to an antigen-specific antibody^15–21^. These systems are designed such that a patient would be infused with both a tagged, antigen-specific antibody that binds to target cells and CAR T cells that react with the tagged antibody on the surface of target cells. Adaptor CARs are also referred to as “universal CARs” as they have the potential to allow for one population of T cells to target multiple tumor antigens by administering different antibodies sequentially or simultaneously. Additionally, the activity of the adaptor CARs can be tuned by altering the concentration of tagged antibodies or halting antibody administration for better control over potential toxicities resulting from over-active CAR T cells. Adaptor CAR systems have been developed recognizing a variety of peptides or small molecules conjugated to antibodies including biotin, fluorescein isothiocyanate (FITC), peptide neo-epitopes (PNE), Fcγ, and leucine zippers, and the first adaptor CAR system is currently in clinical trials^15–21^.

Here, we describe key advances in antigen receptor design – the creation of a switchable adaptor synNotch system and the creation of a novel universal CAR system that acts through self-labeling enzyme chemistry. Our first attempt to create a switchable adaptor synNotch system that functioned through transient binding of the receptor to an antibody was unsuccessful, and we reasoned that a stronger antibody/receptor interaction would be necessary. Seeking to create a self-labeling synNotch receptor that would perform a chemical reaction to covalently fuse with the adaptor antibody, we generated a synNotch receptor containing the SNAPtag protein. SNAPtag is a modified human O-6-methylguanine-DNA methyltransferase (*MGMT*) that was engineered to react to benzylguanine, a bio-orthogonal tag molecule, and is known to be specific and efficient at self-labeling in both cells and animals (**Fig. 1a**)^22–24^. As previous literature for adaptor CARs has shown a positive correlation between CAR function and the CAR/adaptor binding affinity, we also created and characterized a CAR containing the SNAPtag^15,18^. The SNAP-CAR and SNAP-synNotch systems are highly modular receptor platforms for diverse programming of cell behaviors using covalent chemistry (**Fig. 1b,c,d**).

**Fig. 1.**
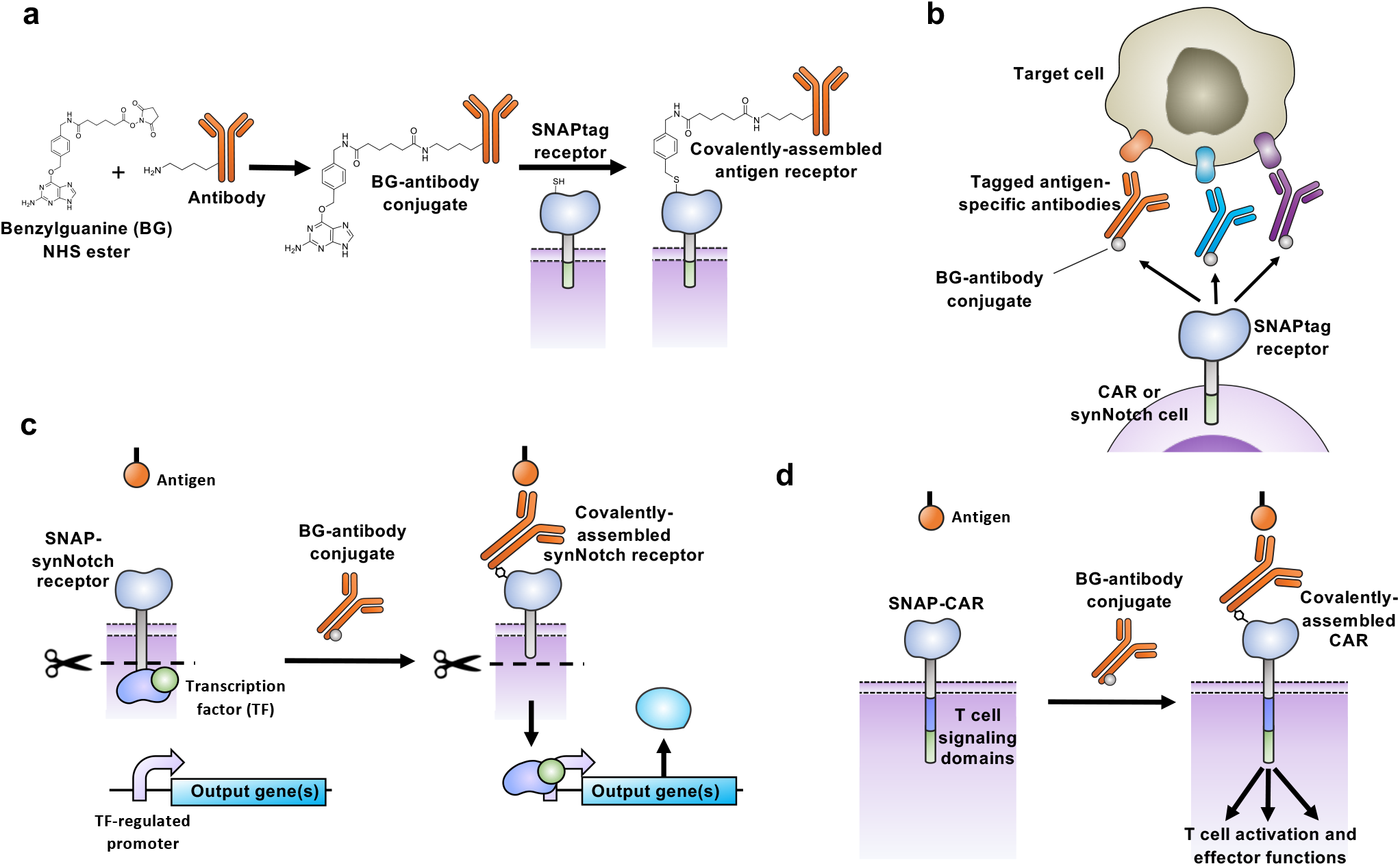
Schematic of switchable adaptor SNAP-CAR and SNAP-synNotch receptor function. **a**, A benzylguanine motif (BG) is chemically-conjugated to an antibody using the benzylguanine NHS ester. The BG-antibody conjugate then fuses to the extracellular SNAPtag enzyme through a self-labeling reaction. **b**, SNAPtag receptors enable the targeting of multiple different antigens using the same receptor by combining SNAP receptor cells with different BG-conjugated antibodies. **c**, The SNAP-synNotch receptor is targeted by a BG-conjugated antibody and upon antigen recognition leads to cleavage of the synNotch receptor, releasing the transcription factor and transcriptional regulation of a target gene or genes. **d**, The SNAP-CAR is targeted by a BG-conjugated antibody to activate T cell signaling and effector functions upon antigen recognition.

## Results

### Engineering a self-labeling SNAP switchable adaptor synNotch receptor

We first created and tested a putative universal synNotch system with a biotin-avidin tag-receptor interaction using the biotin-binding protein, ‘mSA2’ as the targeting domain. The goal of this system was to target receptor specificity by combining mSA2-synNotch cells with biotinylated antibodies, similar to our previously reported mSA2 CAR T cell system ^17^. We cloned the mSA2 synNotch receptor into a lentiviral expression vector and transduced Jurkat cells with the receptor along with a Gal4-driven TagBFP reporter gene (**Supplementary Fig. 1a**). In brief, the process of synNotch receptor activation culminates in the release of the Gal4-VP64 transcription factor from the membrane and transcriptional activation of the Gal4 target gene, for which we are using the TagBFP reporter. Assaying the cells by flow cytometry, we found that the receptor was efficiently expressed on the cell surface (**Supplementary Fig. 1b**). Furthermore, incubating mSA2-synNotch cells with plate-immobilized biotin showed TagBFP response gene activation (**Supplementary Fig. 1c**). However, the receptor was ultimately not functional at detecting cell-surface antigen, as we saw no receptor activation when we incubated the cells with biotinylated antibody-labeled tumor cells (**Supplementary Fig. 1d**). We posited that the lack of signaling by the mSA2 synNotch receptor, in contrast to potent signaling by mSA2 CAR T cells, was the result of the Notch receptor’s differing signaling mechanism that requires a pulling force^25^. We reasoned that a stronger receptor to tag binding interaction (mSA2-biotin K_d_ = 5.5×10^−9^ M) may be required to create a functional, universally applicable synNotch system^26^.

Thus, we next created a synNotch receptor containing the SNAPtag self-labeling enzyme which forms a covalent bond via a benzene ring with a BG-tagged molecule (**Fig. 2a**)^23,24,27^. The goal of this system was to direct receptor activation by combining SNAP-synNotch cells with BG-labeled antibodies (**Fig. 1b,c**). We generated a lentiviral vector encoding the SNAP-synNotch receptor and transduced Jurkat cells (**Fig. 2b**). Antibody labeling of the myc epitope tag and labeling with a fluorophore-conjugated BG reagent confirmed cell surface expression of the receptor and SNAP-BG cell-surface labeling activity (**Fig. 2c**).

**Fig. 2.**
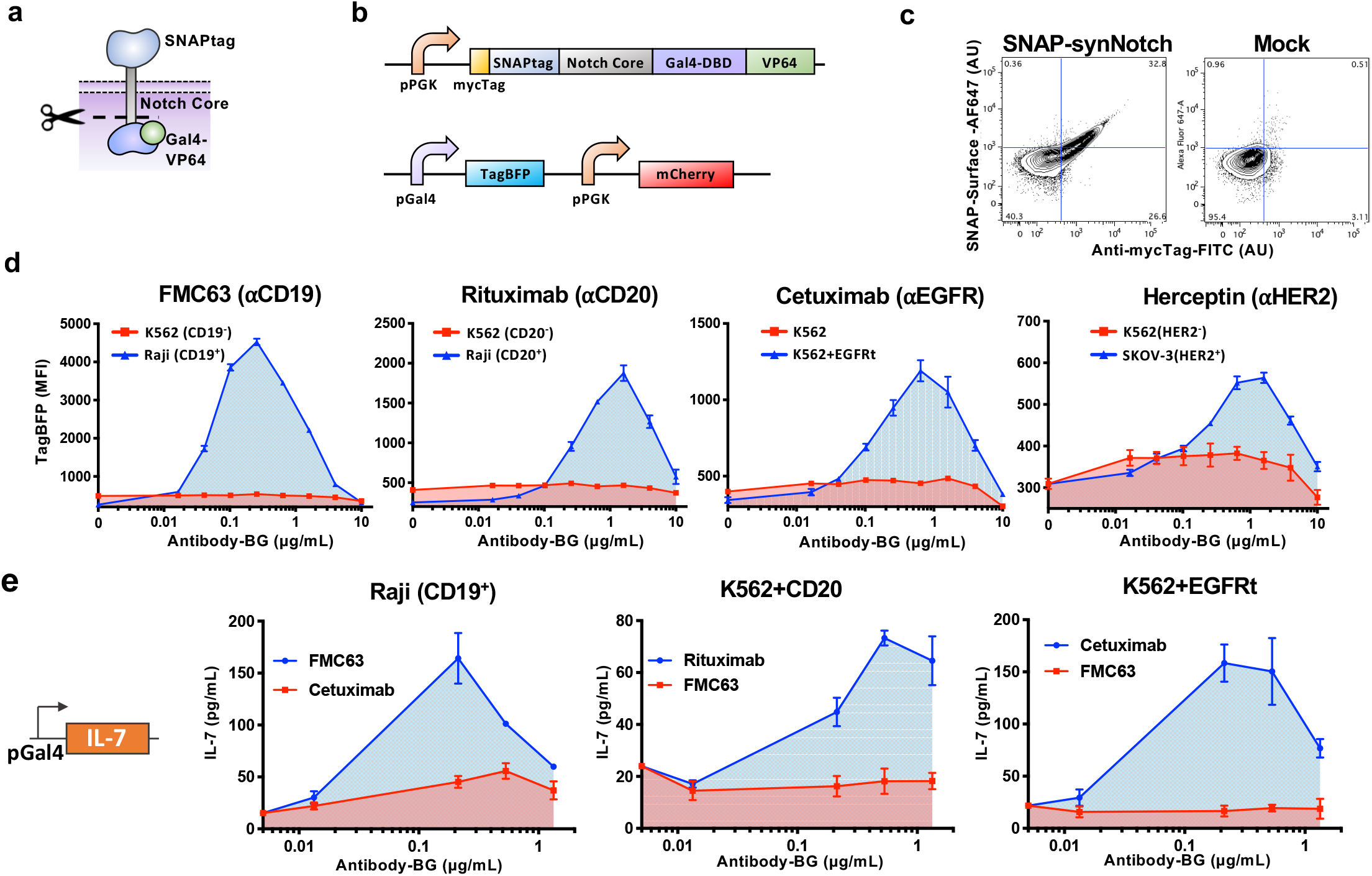
The SNAP-synNotch receptor can be targeted to desired antigens of interest by benzylguanine-conjugated antibodies. **a**, Diagram of the SNAP-synNotch receptor. **b**, Design of SNAP-synNotch receptor expression and response lentiviral vectors. The SNAP synNotch receptor contains the Gal4-VP64 transcription factor which upon activation leads to activation of the TagBFP response gene. **c**, Flow cytometry analysis of surface expression and enzymatic functionality of the SNAP-synNotch receptor on transduced vs. mock (un)transduced Jurkat cells assessed by staining with the anti-Myc-Tag antibody and SNAP-surface-AF647 dye. **d**, Flow cytometry analysis of the activation of SNAP-synNotch cells co-incubated with the indicated target cell lines and antibody concentrations for TagBFP output gene expression reported as mean fluorescence intensity (MFI) gated on mCherry+ cells and **e**, by ELISA for the production of the IL-7 therapeutic transgene. For **d** and **e**, n = 3 biologically-independent experiments ± s.e.m.

To generate the adaptor antibodies, we conjugated BG to lysines or N-termini of several clinically-relevant antibodies using a synthetic BG-NHS ester (**Fig. 1a**). These antibodies included Rituximab targeting CD20, FMC63 targeting CD19, Herceptin targeting HER2, and Cetuximab targeting EGFR^28–31^. While the conjugation products were heterogenous, we quantified the average number of BG molecules conjugated to each antibody by a SNAP protein-labeling assay in which SNAP-conjugation led to a shift in the antibodies’ molecular weight that could be resolved by SDS-PAGE (**Supplementary Fig. S2**). The frequency of BG molecules per antibody ranged from 2.0-2.8 as summarized in **Supplementary Table S1**. We characterized the antigen expression and BG-antibody staining of various target cell lines by flow cytometry in which the antibodies displayed expected antigen specificities (**Supplementary Fig. S3**).

Next, we tested the BG-conjugated FMC63 antibody (FMC63-BG) for its ability to activate synNotch signaling in response to CD19 positive tumor cells. We performed a co-incubation assay of SNAP-synNotch cells and CD19 positive and negative tumor cells in the presence of different levels of the FMC63-BG antibody conjugate, and after 48 hours we assayed for TagBFP response gene expression by flow cytometry. TagBFP expression was significantly up-regulated in response to CD19 positive tumor cells for various concentrations of antibody. Receptor activation was sensitive, with significant activation observed at an antibody concentration as low as 0.04 μg/mL and increasing to a peak at 0.25 μg/mL. Response gene activation then decreased with increasing antibody amounts before being completely inhibited at a concentration of 10 μg/mL, indicative of a “hook effect”, in which case the antibody is in such excess that different antibody molecules are saturating both the target cells and synNotch cells without formation of ternary complexes. This behavior is commonly observed with chemical and cell processes that involve ternary complex formation (**Fig. 2d**) ^32^.

We found that BG-conjugated antibodies targeting other antigens were also capable of activating the SNAP-synNotch receptor in an antigen-specific manner (**Fig. 2d**). We performed similar co-incubation assays of SNAP-synNotch cells and antigen positive and negative tumor cells in the presence of different levels of BG-conjugated Cetuximab, Herceptin, and Rituximab antibodies and assayed for TagBFP response gene expression by flow cytometry. Significant up-regulation of TagBFP was again observed for each of the tested antibodies, tunable through increasing antibody concentrations, and in a target antigen-specific manner. A similar activation pattern was observed for each of the antibodies – an increase with the antibody dose until a peak activation level followed by an antibody dose-dependent decrease. Receptor activation was also target cell dose-dependent, having optimal activity at high target to SNAP-synNotch cell ratios (**Supplementary Fig. S4**).

Next, we tested whether SNAP-synNotch receptor was capable of regulating expression of the IL-7 response gene, a candidate therapeutic gene of interest for its ability to promote T cell proliferation^33^. We generated an IL-7 response gene expression construct in which the TagBFP gene was replaced by the IL-7 coding region and placed under synNotch control^33^. This construct was again transcriptionally activated by the Gal4-VP64 transcription factor upon receptor activation. We transduced SNAP-synNotch cells with this response vector and co-incubated them with several different antibodies and antigen positive and negative tumor cells for evaluation of IL-7 response gene expression by ELISA (**Fig. 2e**). Similar to TagBFP response gene activation, IL-7 was significantly up-regulated in an antigen-specific and antibody dose-responsive manner demonstrating the modularity of the SNAP-synNotch system in controlling different output genes.

### Engineering a self-labeling SNAP switchable adaptor CAR

We next aimed to create a switchable adaptor CAR system using the SNAPtag protein domain to target T cell receptor signaling when combined with BG-tagged antibodies (**Fig. 1d**). We cloned the SNAPtag domain into a lentiviral vector containing the standard CAR components including the CD8⍰. hinge and transmembrane domains, the 4-1BB cytoplasmic co-signaling domain, the CD3zeta T cell signaling cytoplasmic domain, and a TagBFP reporter gene co-expressed via a T2A co-translational peptide (**Fig. 3a,b**). We packaged this vector into lentivirus particles and transduced Jurkat cells. We found that the receptor was efficiently expressed and that the SNAPtag protein was functional as indicated by TagBFP expression and staining with a BG-fluorophore reagent and analysis by flow cytometry (**Fig. 3c**).

**Fig. 3.**
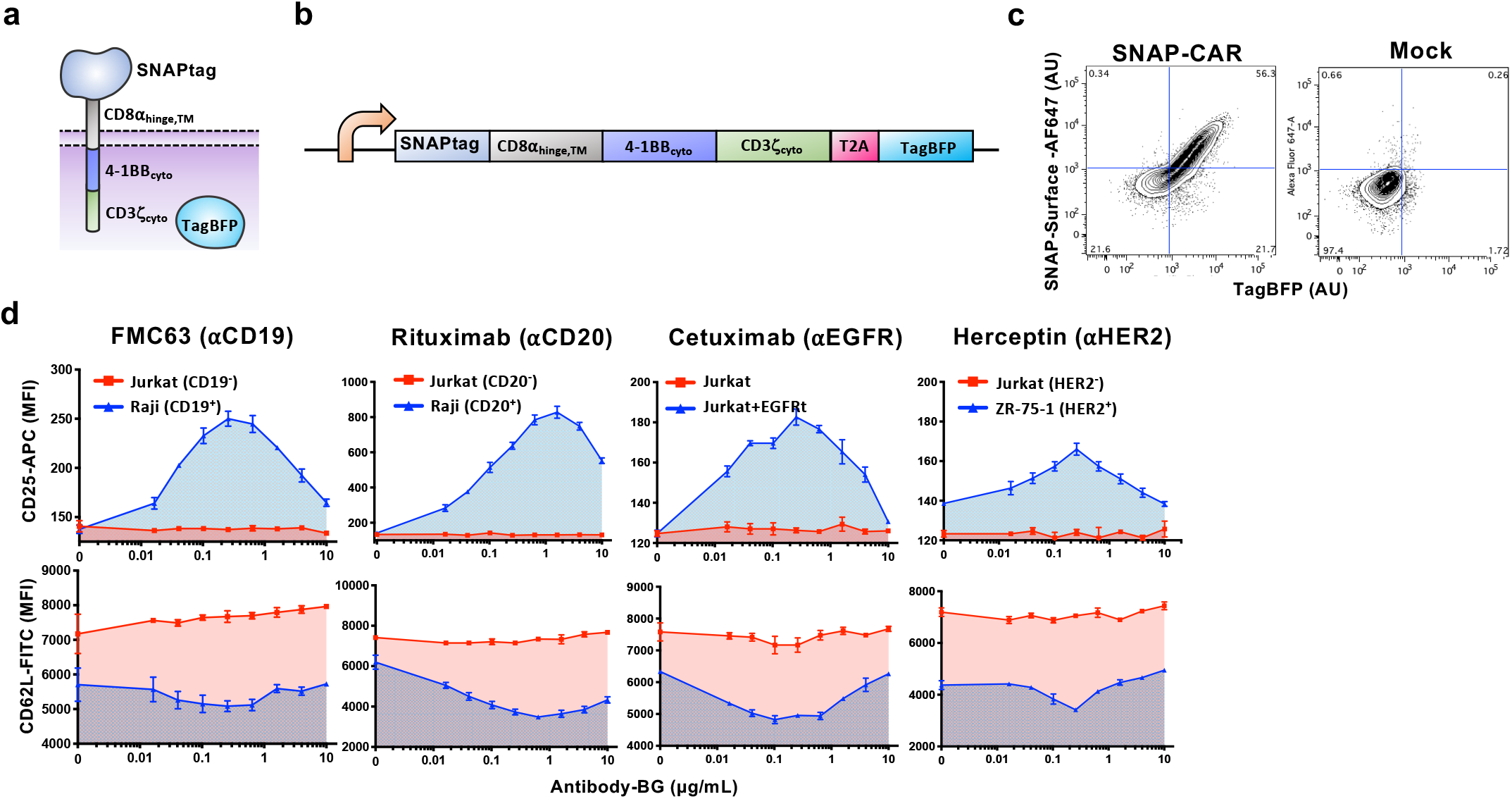
The SNAP-CAR can be targeted to desired antigens of interest by benzylguanine-conjugated binding proteins. **a**, Diagram of the SNAP-CAR. **b**, Design of the SNAP-CAR lentiviral expression construct. **c**, Flow cytometry analysis of the expression and enzymatic functionality of the SNAP-CAR receptor on transduced vs. mock (un)transduced Jurkat cells assessed by staining with SNAP-surface-AF647 dye and recording TagBFP expression. **d**, Flow cytometry analysis of CD25 and CD62L T cell activation markers on Jurkat SNAP-CAR effector cells co-incubated with the indicated target cell lines and antibody concentrations reported as mean fluorescence intensity (MFI). CD25 increases while CD62L decreases with activation, n = 3 biologically-independent experiments ± s.e.m.

We next tested whether BG-conjugated antibodies could be combined with SNAP-CAR Jurkat cells to target their T cell activation signaling. We co-incubated SNAP-CAR cells with various antigen positive or negative tumor cell lines and various doses of BG-conjugated antibodies. After 24 hours we assayed the cells for T cell activation by staining with antibodies specific for CD25, which is up-regulated upon T cell activation, and CD62L, which is down-regulated. We assessed the expression levels of these markers in SNAP-CAR cells by specifically gating on the TagBFP CAR+ cell population. We found that expression of these markers was controlled in an antigen-specific and dose-responsive manner by the BG-conjugated antibodies (**Fig. 3d**). Similar to SNAP-synNotch cells, SNAP-CAR activation signaling strength peaked at an antibody dose level between 0.1-1.0 μg/mL before steadily decreasing, again indicative of a hook effect.

### The SNAP-CAR is functional in primary human T cells

We then tested the expression level and *in vitro* functional activity of the SNAP-CAR in primary human T cells transduced with the SNAP-CAR lentivirus. Staining with a BG-fluorophore conjugate and assaying by flow cytometry, we found that the SNAP receptor was efficiently expressed in ~40% of cells in a manner that correlated well with the expression of the TagBFP marker gene (**Fig. 4a**). To test CAR functionality, we co-incubated SNAP-CAR T cells with various antigen positive or negative target tumor cell lines and 1.0 μg/mL of BG-conjugated antibodies for 24 hours. Targeted antigens included CD20, EGFR, and HER2. Analyzing the supernatants of the co-incubated cells by ELISA we found that the SNAP-CAR T cells could be directed by the covalently attached BG-antibodies to produce significant amounts of IFNγ in response to antigen positive target cells (**Fig. 4b**). Our analysis of co-incubated cells by flow cytometry revealed that the antibodies also led to induction of T cell activation markers, up-regulation of CD69 and CD107a, and down-regulation of CD62L (**Fig. 4c**). CAR cells and target cells were also co-cultured and evaluated for target cell lysis by flow cytometry and showed high levels of target specific cell lysis (**Fig. 4d**). Again, T cell marker activation and target cell lysis were significantly higher only when the co-administered antibody targeted an antigen expressed by the co-administered cells, indicating antibody specificity. In addition to full length IgG antibodies, we also tested a BG-conjugated Fab fragment of Rituximab. This molecule, more similar to the scFv antibody fragment found in traditional CARs, also showed potent activity for each of the effector functions equal to or greater than that of the full-length Rituximab.

**Fig. 4.**
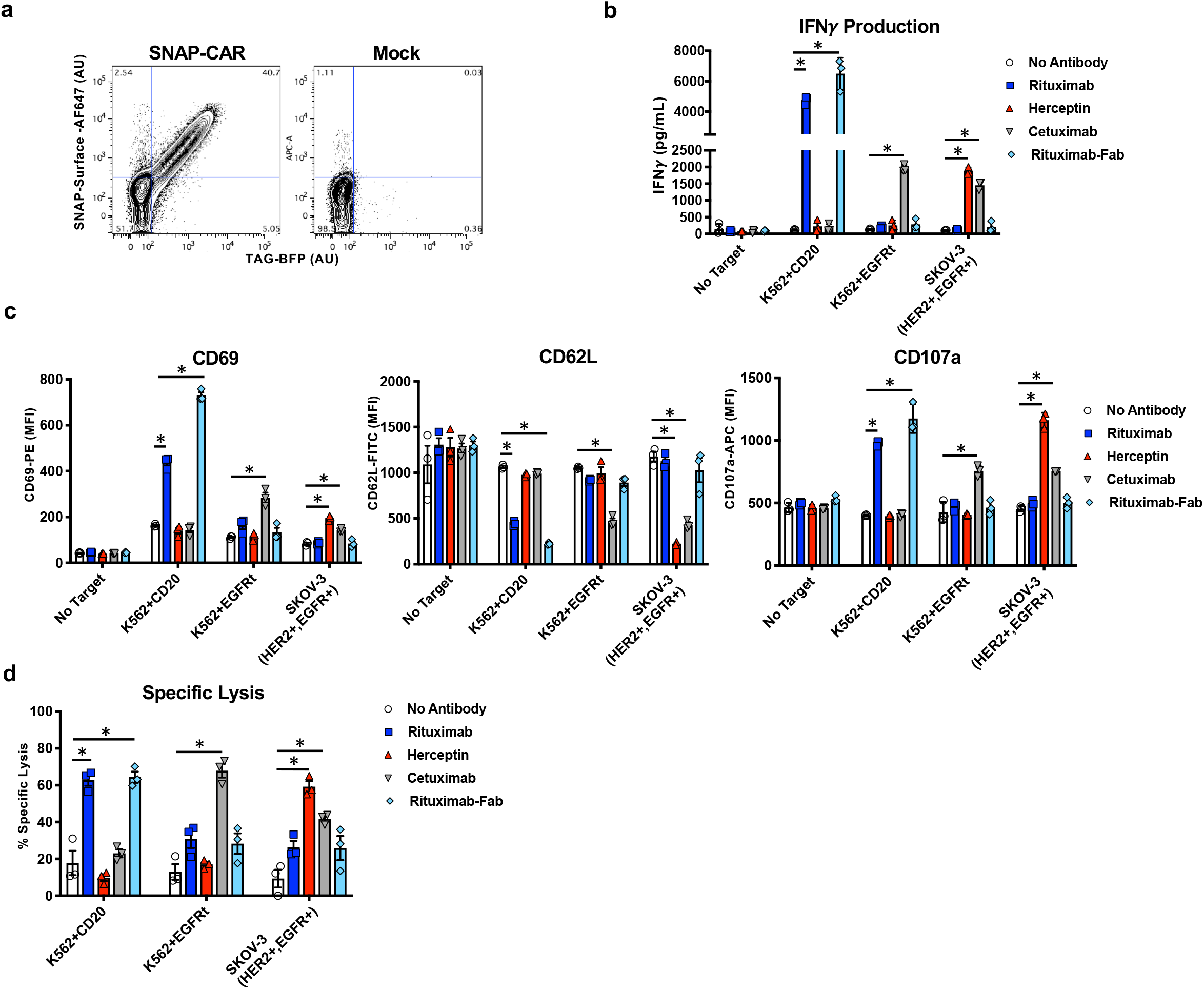
The SNAP-CAR is effective on primary human T cells. **a**, Flow cytometry analysis of the expression and enzymatic functionality of the SNAP-CAR on transduced vs. mock (un)transduced primary human T cells by staining with SNAP-surface-AF647 dye and recording TagBFP expression. **b**, ELISA for IFNγ production from primary human SNAP-CAR T effector cells co-incubated with the indicated target cell lines and 1.0 μg/mL of the indicated antibody and **c**, flow cytometry analysis of CD69, CD62L, and CD107a T cell activation markers on the SNAP-CAR (TagBFP+) population from the co-incubations in *b*, reported as MFI. **d**, Specific lysis of target cell lines by co-incubated primary human SNAP-CAR T cells and 1.0 μg/mL of the indicated BG-conjugated antibodies. For **b, c** and **d**, multiple ANOVA comparisons were performed. As the data did not have homogeneity of variance (Levene’s test), Tukey’s HSD was used for post-hoc analysis between antibody conditions. “*” denotes a significance of p < .0001, n = 3 biologically-independent experiments ± s.e.m.

### BG-antibody pre-loading experiments to interrogate the receptor signaling mechanism

To further investigate the mechanism of SNAP-synNotch receptor activation, we performed co-incubation assays in which we pre-labeled either the target cells or SNAP-synNotch cells with BG-conjugated antibodies. We incubated CD19 positive and negative tumor cells with BG-conjugated antibody, washed away residual unbound antibody, and co-incubated these cells with SNAP-synNotch cells for 48 hours. Evaluating response gene activation, we found that CD19 positive tumor cells significantly up-regulated TagBFP gene expression to a level comparable to the peak level of activation in the previous dose-response assay (**Fig. 5a, 2c**). We then pre-labeled the SNAP-synNotch cells with BG-conjugated antibody and after washing away residual antibody, we performed a similar co-incubation assay with CD19 positive or negative tumor cells. No significant response gene activation was observed (**Fig. 5b**).

**Fig. 5.**
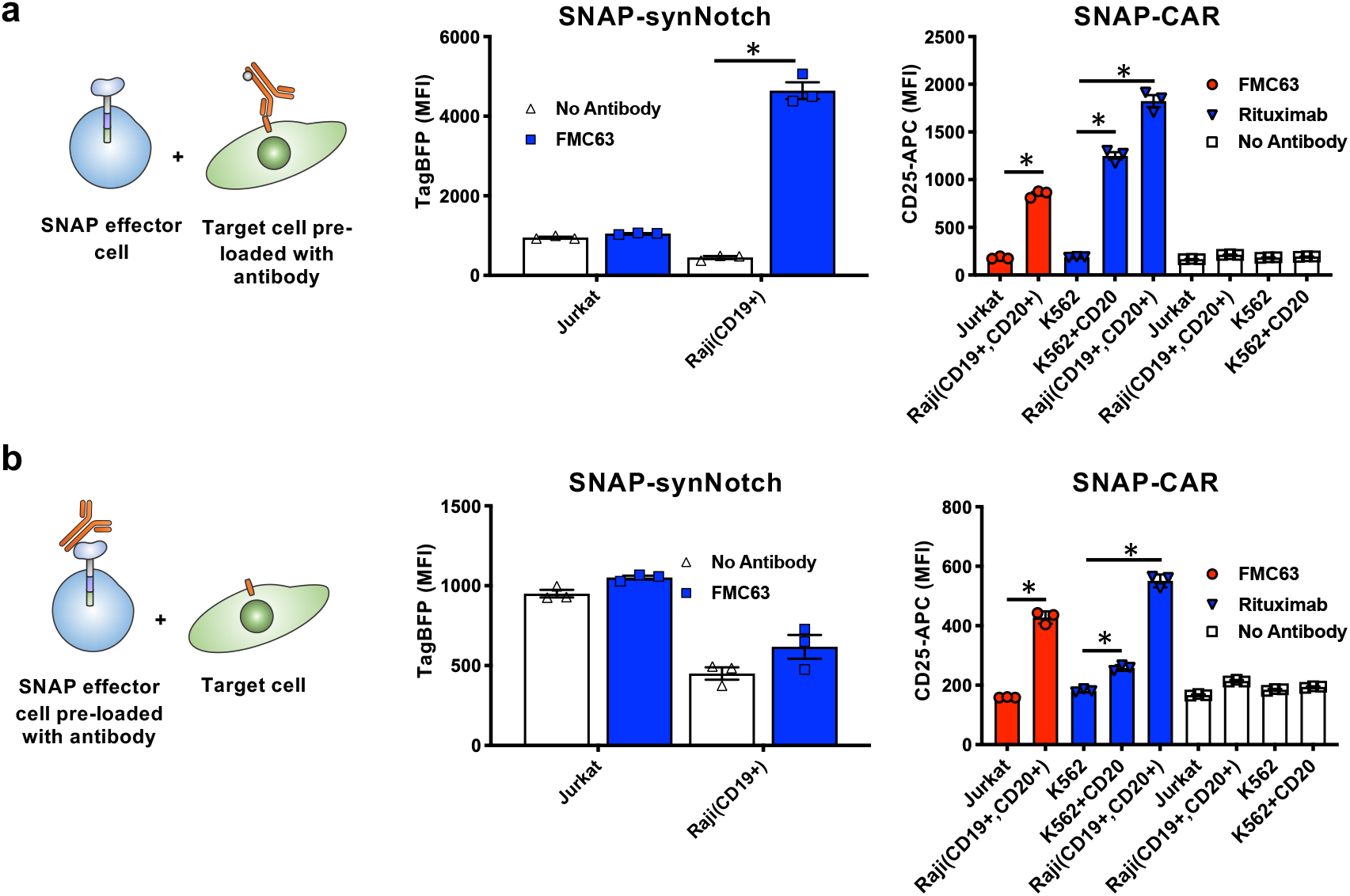
Characterizing the activity of SNAP receptors when pre-assembled or with pre-labeled target cells. **a**, Flow cytometry analysis of SNAP receptor activation for SNAP-synNotch and SNAP-CAR cells co-incubated with target cells that were pre-labeled with the indicated antibodies. **b**, Flow cytometry analysis of SNAP receptor activation for SNAP-synNotch and SNAP-CAR cells that were pre-labeled with the indicated antibodies and co-incubated with target cells. MFI of TagBFP output gene expression and CD25 marker expression were evaluated by flow cytometry for SNAP-synNotch and SNAP-CAR cells, respectively. For **a** and **b**, multiple ANOVA comparisons were performed. As the data did not have homogeneity of variance (Levene’s test), Tukey’s HSD was used for post-hoc analysis between antibody conditions. “*” denotes a significance of p < .0001, n = 3 biologically-independent experiments ± s.e.m.

Performing the same pre-staining experiments with SNAP-CAR cells, we found that the SNAP-CAR was functional both when the SNAP-CAR cells or tumor cells were pre-labeled with BG-conjugated antibodies. We incubated antigen positive or negative tumor cells with BG-conjugated antibody, washed away residual unbound antibody, and co-incubated these cells with Jurkat SNAP-CAR cells for 24 hours. Assaying by flow cytometry for the CD25 T cell activation marker, we found that the labeled tumor cells induced a significant up-regulation of CD25 expression to a level comparable to the peak level of activation in the previous dose-response assay (**Fig. 5a**). We then pre-labeled the SNAP-CAR cells with BG-labeled antibody and after washing away residual antibody, we performed a similar co-incubation assay with CD20 or CD19 positive and negative tumor cells. Significant up-regulation of T cell activation was observed, however, to a lower magnitude than that with pre-labeled cancer cells (**Fig. 5b**).

### Mathematical model of switchable adaptor complex formation

To gain a better understanding of the switchable adaptor receptor signaling and the observed hook effect, we generated a continuous mathematical model of the ternary complex formation between T cell, adaptor antibody, and target cell. Using Python Jupyter Notebook, we created a generalizable model of ordinary differential equations (ODEs) that describe the binding reactions. A system of equations was defined to describe the accumulation and concentration of each of the six species in the model: T cells, antibodies, tumor cells, T cells bound to antibody, tumor cells bound to antibody, and T cell-antibody-tumor cell ternary complexes (**Fig. 6a, Methods**).

**Fig. 6.**
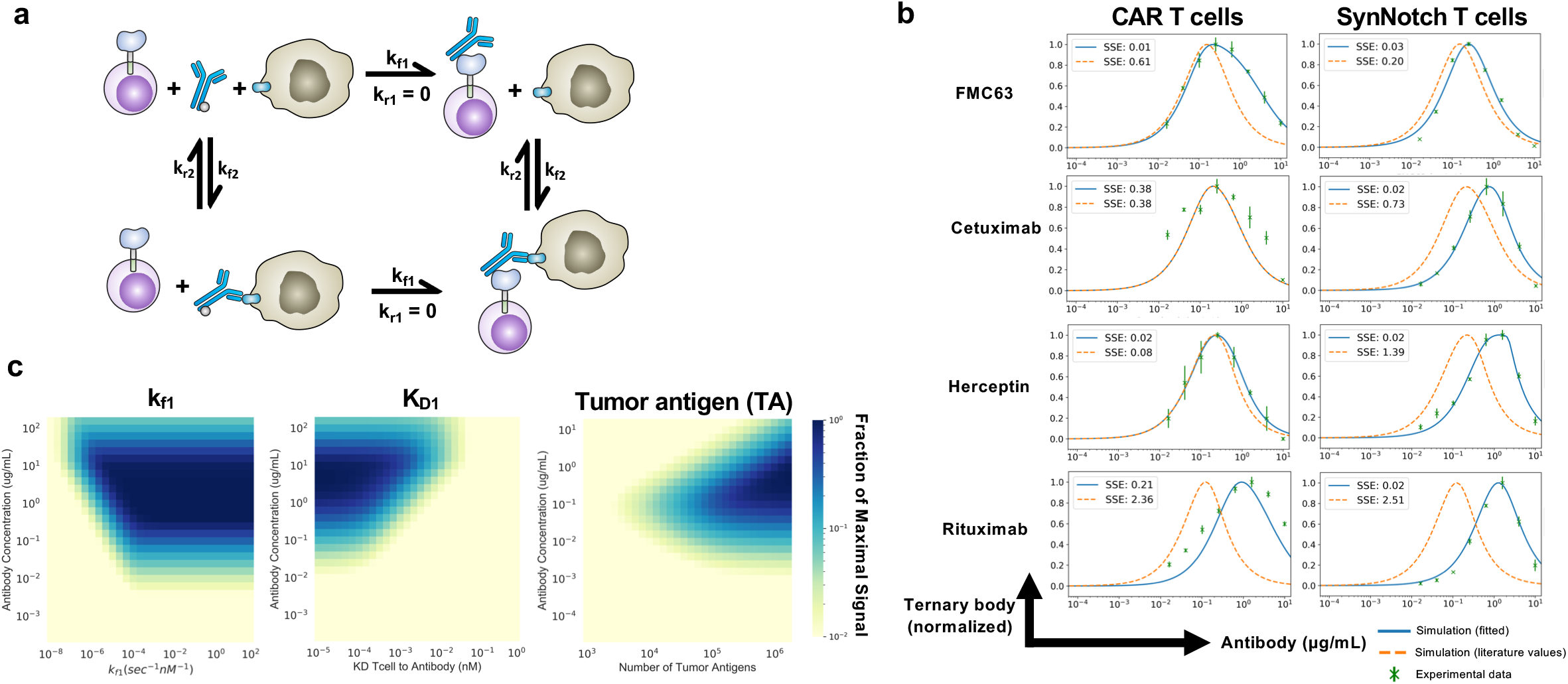
Mathematical model of three-body binding in the context of antibody mediated T cell targeting. **a,** Schematic of the ODE model for SNAP receptor ternary body formation. **b,** Model simulations using parameters from the literature and from parameter estimation, compared to experimental results for four different antibody and antigen pairs for SNAP-CAR and SNAP-synNotch receptors. **c,** Parameter scans of k_f1_ (binding rate of T cells to antibody), K_D1_ (equilibrium dissociation constant between T cell and antibody), and the number of target antigens on the surface of the tumor.

To validate the model, we ran simulations using kinetic parameters taken directly from the literature and then through bounded parameter fitting (**Supplementary Table S2**)^34–37^. Using direct literature values, the model was able to recapitulate the general features of our experimental data, including the observed hook effect and a prediction accurate within an order of magnitude of each antibody dose expected to yield maximum receptor signaling (**Fig. 6b**). We used the sum of squared error (SSE) calculations to measure error in model simulations against experimental results. With the exception of Rituximab, model simulations using literature values alone, resulted in good recapitulations of experimental data (average SSE with literature values = 1.03) (**Supplementary Table S3**). Next, using SciPy, we minimized the sum of squared error to optimize the kinetic parameters and better fit the model to the experimental data for each antibody pair^38^. During parameter estimation, the literature values were used as initial estimates and bounded within one order of magnitude. With these constraints, we were able to minimize our model error in seven of the experimental results to (average SSE after fitting = 0.09) (**Supplementary Table S3**).

With a validated model, we next aimed to use the model to predict how different system parameters would affect receptor signaling and conducted parameter scans with the model. We first varied k_f1_, the forward reaction rate of the antibody binding to the T cell receptor, to simulate the effects increasing the on-rate, which could also be experimentally varied by changing the number of BG’s conjugated to an antibody. We found that the model predicted that increasing the number of BG’s per antibody would lead to greater ternary body formation at higher concentrations of antibody. Once the k_f1_ rate becomes greater than a threshold of 10^−3^ nm^−1^sec^−1^, this effect was expected to plateau. (**Fig. 6c**). Next, we scanned different values for the antibody to T cell affinity, the parameter maximized by the use of the covalent SNAP to antibody interaction. We found that stronger affinity was predicted to lead to ternary body formation over a wider range of antibody concentrations and to lead to a higher overall level of ternary body formation. Finally, as target antigen concentration can vary based on the antigen being targeted, and for cancer, expression levels can also significantly vary greatly between patients and on cells within the same patient, we performed a parameter scan varying the level of tumor antigen concentration. We found that greater antigen concentrations were expected to broaden the effective antibody dosage window for successful ternary complex formation, while lower antigen levels were predicted to require a higher amount of antibody to induce signaling, and were predicted to be more susceptible to inhibition by the hook effect.

## Discussion

The switchable adaptor SNAP-synNotch system further increases the versatility of the synNotch receptor framework leading to post-translational control of receptor signaling by co-administered antibody dose, as well as the ability to target multiple antigens using a single genetically-encoded receptor. While other adaptor CARs have been described, this is, to our knowledge, the first adaptor synNotch system. Our unsuccessful initial attempts to create an adaptor synNotch system using a non-covalent interaction (between mSA2 and biotin) suggest that a very high affinity interaction (ideally covalent) between the synNotch receptor and the tag would be required for the adaptor system to function; presumably since the signaling mechanism for Notch is based on an actual pulling force^25^.

With the covalent bond generated by the self-labeling enzyme, the SNAP-CAR has several beneficial characteristics over other adaptor CAR technologies^15–21^. The results of others, and our results supported by our modeling analysis, suggest that the affinity of the interaction between the CAR and the adaptor molecule is a key parameter for productive receptor signaling^15,18^. The covalent bond produced by the SNAP enzyme and the benzylguanine moiety provides the tightest theoretical bond – a covalent bond – and thus maximizes this critical parameter. While many antibodies will be functional with a non-covalent, lower-affinity adaptor CAR, our model predicts that covalent bond formation could enable use of antibodies that otherwise have a binding affinity to the target antigen that is too low to elicit an effect. Furthermore, the SNAP enzyme reacting to the bio-orthogonal benzylguanine grants the CAR exquisite specificity, and, being an enzyme of human origin, the SNAP protein is likely to be well-tolerated in a human host. This characteristic satisfies a key requirement for the persistence of the adoptively transferred therapeutic cells and minimizes the possibility of toxicities resulting from their immune rejection^39–41^.

The ability to create functional CARs by preloading the SNAP receptor, followed by removal of excess BG-antibody, provides unique opportunities to test candidate antigen binding regions as components of traditional CARs (**Fig 5b**). Compared to adaptor CARs binding to antigen through a transient interaction, the covalently assembled receptor will more closely resemble that of a traditional CAR. We were originally surprised that the pre-assembled SNAP-synNotch receptors were not functional and instead require pre-targeting of the cancer cells. However, this result can potentially be explained by considering the mechanism of signaling, in which the receptor is proteolytically cleaved and thus destroyed following activation, not allowing for multiple signaling events from recycled receptors. This result suggests that multiple bursts of receptor activation from distinct receptors over time may be needed to sufficiently trigger synNotch signaling. As the pre-assembled CARs were capable of signaling, pre-loading the SNAP-CAR T cells could be a potential clinical approach, however, upon T cell activation, the cells would be induced to expand thus diluting out the assembled receptor, requiring supplementation of additional antibody. Similarly, while the SNAP-CAR T cells showed significant up-regulation of T cell activation markers for both pre-loading conditions, more robust T cell activation was observed with pre-loaded tumor cells than pre-loaded SNAP-CAR cells. That both the CAR and SynNotch receptors were maximally effective against tumor cells pre-labeled with antibodies suggests that pre-dosing of a patient with tagged antibody prior to T cell infusion could be the optimal treatment regimen.

Based on our results, additional self-labeling or covalent protein assembly systems could also provide good frameworks for universal adaptor CARs. Such systems include candidates such as: CLIPtag, Halotag, SpyTag, SnoopTag, Isopeptag, Sortase or split inteins ^23,42–45^. It is possible that CARs made with other self-labeling enzymes, having different kinetic binding/dissociation rates, molecular size, and could vary in expression level, could provide more robust signaling depending on the characteristics of the antigen(s) being targeted. Additionally, other synthetic receptor platforms could be amenable to the switchable adaptor format by creating receptors with the SNAPtag protein domain^46,47^.

Future developments of the SNAP adaptor systems to clinical applications will be important in several key areas. Developing site-specific tagging approaches will help to lead to more homogeneous antibody-BG conjugates and potentially identify optimal BG-conjugation sites that could further maximize receptor signaling output. For any specific disease indication there is the need to determine the optimal antibody or antibodies, to combine with the adaptor T cells to provide disease-specific or at least disease-associated induction of receptor signaling. While it is known that SNAP self-labeling is efficient in animal models, testing the SNAP T cells with specific adaptor antibody combinations will be important to confirm potent function and to optimize the therapeutic delivery strategy^22^.

Our molecular model of switchable receptor systems provided key insights into our observed signaling behaviors and yielded predictions on how to potentially optimize receptor function. The model results suggested that the binding strength between the CAR and the adaptor is a critical parameter for signaling and that our SNAP receptors for which this interaction strength is maximized via a covalent bond are expected to be desirable. Furthermore, the model suggested that one way to improve activity would be to increase the forward reaction rate of the CAR binding to the adaptor, which could potentially be accomplished by increasing the number of BG molecules per antibody. Lastly the model predicted that using adaptor CARs to target antigens that are expressed at high levels is preferable as these antigens would be expected to induce receptor signaling at lower antibody concentrations and would be less susceptible to the hook effect at higher antibody doses.

SNAP-synNotch and SNAP-CAR T cells provide a powerful new adaptor strategy for fully programmable targeting of engineered cells to multiple antigens using covalent chemistry. These systems have great potential for clinical application and biotechnological utility by providing researchers with the ability to rapidly screen CAR and synNotch antibody candidates and to rewire and activate cellular programs in response to highly specific antibody-antigen interactions.

## Methods

### Construction of lentiviral expression vectors

pHR_PGK_antiCD19_synNotch_Gal4VP64 and pHR_Gal4UAS_tBFP_PGK_mCherry were gifts from Wendell Lim (Addgene plasmid# 79125; http://n2t.net/addgene:79125; RRID:Addgene_79125 and Addgene plasmid# 79130; http://n2t.net/addgene:79130; RRID:Addgene_79130, respectively). Sequences for all receptor coding regions and response constructs are listed in **Supplementary Table S4**. To generate pHR-PGK-SNAP-41BBζ, a DNA fragment encoding SNAP-41BBζ was codon-optimized, synthesized (Integrated DNA Technologies) and cloned into the pHR-PGK vector backbone using isothermal assembly. To generate pHR-PGK-SNAP-synNotch-Gal4VP64 and pHR-PGK-mSA2-synNotch-Gal4VP64, DNA encoding the SNAP or mSA2 coding region was codon-optimized and synthesized (Integrated DNA Technologies) and cloned in place of the anti-CD19scFv in plasmid pHR-PGK-antiCD19-synNotch-Gal4VP64 (Addgene# 79125) using isothermal assembly. To generate pHR-Gal4UAS-IL7-PGK-mCherry, a DNA fragment encoding IL-7 was codon-optimized, synthesized (Integrated DNA Technologies) and cloned in place of TagBFP in the pHR_Gal4UAS_tBFP_PGK_mCherry vector backbone using isothermal assembly. Virus was generated using the above described transfer vectors following methods described previously in detail^48^.

### Production of BG-antibody conjugates

Rituximab (Rituxan, Genentech), Cetuximab (Erbitux, Eli Lily), and Herceptin (Traztuzumab, Genentech) and FMC63 (Novus Biologicals) underwent buffer exchange into PBS using 2 mL 7K MWCO Zeba Spin Desalting Columns (ThermoFisher Scientific). The Rituximab Fab fragment was generated using the Fab Preparation Kit (Pierce) following the manufacturer’s protocol. Antibodies were then co-incubated with a 20-fold molar excess of BG-GLA-NHS (NEB) for 30 minutes at room temperature, followed by buffer exchange into PBS using 2 mL 7K MWCO Zeba Spin Desalting Columns.

### Quantification of BGs on BG-conjugated antibodies

For *in vitro* conjugation of whole antibodies with SNAPtag, BG-conjugated purified antibodies (0.5 μg) were incubated with recombinant SNAPtag protein (2 μg). The solution was incubated in PBS (10 μL, pH 7.4) containing DTT (1 mM) at 37°C for two hours. Conjugation solutions were then diluted with Laemmli buffer, boiled for 5 minutes, and analyzed on an 8% SDS-PAGE (120 V, 1.5 h). Gels were visualized using imidazole-SDS-Zn reverse staining.1 Briefly, gels were stained with a 200 mM imidazole aqueous solution containing 0.1% SDS for 15 minutes with light agitation. The staining solution was decanted and replaced with water. After 30 seconds, the water was decanted and the gel was developed for 45 seconds with a 200 mM ZnSO4 aqueous solution with light agitation. The gel was then rinsed under running water for 10 seconds. Gels were imaged on a ChemiDoc (Bio-Rad) using epi white light on a black background. Relative band intensities were quantified with ImageJ. A correction factor of 1.5 was applied to the average number of BG/antibody to account for the light chain. Light chains were conjugated to SNAPtag in the same manner except 3 μg of antibody was incubated with 6 μg SNAPtag, gels were analyzed on a 10% SDS-PAGE (120 V, 1.2 h) and stained with Coomassie, and a correction factor was not applied.

### Cell line culture

Human tumor cell lines Jurkat Clone E6-1 (TIB-152), ZR-75-1(CRL-1500), K562 (CCL-243), SKOV-3(HTB-77), and Raji (CCL-86) were obtained from American Type Culture Collection (ATCC) and cultured at 37°C in RPMI medium supplemented with 1X MEM amino acids solution, 10mM Sodium Pyruvate, 10% fetal bovine serum (FBS), and Penicillin-Streptomycin (Life Technologies). K562+EGFRt, K562+CD20, and Jurkat+EGFRt cells that stably express full-length CD20 and the EGFRt gene https://www.ncbi.nlm.nih.gov/pmc/articles/PMC3152493/, were generated by transducing Jurkat cells with the indicated tumor antigen expressing lentivirus and sorting for cells positive for antigen expression. To create the SNAP-CAR stable cell line, Jurkat cells were transduced with SNAP-41BBζ, and underwent fluorescence-activated cell sorting (FACS) for TagBFP expression. and reporter (mCherry+) expression. To generate SNAP-synNotch lines, SNAP-synNotch-Gal4VP64 was co-transduced with either pHR-Gal4UAS-tBFP-PGKmCherry or pHR-Gal4UAS-IL7-PGKmCherry lentivirus, and receptor and response construct positive cells were obtained by FACS for anti-Myc-Tag antibody staining (Cell signaling Technology) and mCherry expression, respectively. HEK293T cells (ATTC, CRL-3216), used for lentivirus production were cultured at 37°C in DMEM supplemented with 10% FBS, and Penicillin-Streptomycin.

### Primary human T cell culture and transduction

All primary T cells for experiments were sourced from deidentified human Buffy Coat samples purchased from the Pittsburgh Central Blood Bank fulfilling the basic exempt criteria 45 CFR 46.101(b)(4) in accordance with the University of Pittsburgh IRB guidelines. PBMC were isolated from a Buffy Coat from healthy volunteer donors using Ficoll gradient centrifugation and human T cells were isolated using the Human Pan T cell isolation kit (Miltenyi Biotec). Human T cells were cultured in supplemented RPMI media as described for cell lines above, however, 10% Human AB serum (Gemini Bio Products) was used instead of FBS, and the media was further supplemented with 100 U/ml human IL-2 IS (Miltenyi Biotec), 1 ng/ml IL-15 (Miltenyi Biotec), and 4mM L-Arginine (Sigma Aldrich). T cells were stimulated and expanded using TransAct Human T cell activation reagent (Miltenyi Biotec). For transduction, 48 hours after activation, lentivirus was added to cells at a multiplicity of infection of 10–50 in the presence of 6 mg/ml of DEAE-dextran (Sigma Aldrich). After 18 hours, cells were washed and resuspended in fresh T cell media containing 100 U/ml IL-2 and 1 ng/ml IL-15. Cells were split to a concentration of 1M/mL and supplemented with fresh IL-2 and IL-15 every 2-3 days. After 10-12 days of stimulation and expansion, transduced cells were evaluated for CAR expression by flow cytometry and evaluated for activity in subsequent functional assays.

### Flow cytometry staining

Cells were washed and resuspended in flow cytometry buffer (PBS + 2% FBS) and then stained using the indicated antibodies for 30 minutes at 4°C followed by two washes with flow cytometry buffer. Live cells and singlets were gated based on scatter. To evaluate SNAP-CAR and SNAP-synNotch expression, 1M cells were labeled with SNAP-Surface 647 (NEB) following manufacturer’s recommendation (50 μM concentration of SNAP-Surface 647 in complete cell media) for 30 minutes at 37°C and washed three times in complete culture media. SNAP-synNotch cells were additionally stained with anti-Myc-Tag-FITC antibody (Cell Signaling Technology) to stain the Myc-Tag on the N-terminus of the receptor.

### SNAP-synNotch cell and target cell co-incubation assays for antibody-mediated activation

100,000 synNotch Jurkat effector cells were co-cultured with 200,000 of the indicated target cells and BG-conjugated antibody for 48 hours. For assays with SNAP-synNotch cells engineered with the pHR_Gal4UAS_tBFP_PGK_mCherry response construct, co-incubated cells were evaluated by flow cytometry, gating for synNotch cells by mCherry positivity, and then quantifying TagBFP fluorescence for this mCherry+ population. For assays with SNAP-synNotch cells engineered with the pHR_Gal4UAS_IL7_PGK_mCherry construct, following a 48 hour co-incubation, cells were spun down and supernatants were collected and analyzed by ELISA for IL-7 following the manufacturer’s recommended protocol (Peprotech).

### SNAP-CAR T cell and target cell co-incubation assays for antibody-mediated activation

100,000 SNAP-CAR Jurkat or primary human T cell effector cells were co-incubated with 200,000 of the indicated target cells and antibody concentrations for 24 hours and assayed by flow cytometry for T cell marker gene expression. For primary cell assays, cells were stained with CD69-PE (BD Biosciences), CD62L-FITC (BD Biosciences), and CD107a-APC (BD Biosciences) antibodies and for Jurkat effector assays, cells were stained with CD62L-FITC (BD Biosciences) and CD25-APC (BD Biosciences) antibodies. For flow cytometry CAR+ cells were analyzed by gating for the TagBFP+ population. Supernatants from primary cell assays were also collected and analyzed for IFN◻ by ELISA (BioLegend). All assays were performed in triplicate and average IFN◻ production was plotted with standard deviation.

### Target cell lysis assay

The indicated target cells were stained with Cell Trace Yellow following the manufacturer’s recommended protocol (ThermoFisher), and 10,000 target cells per well were co-cultured with 50,000 SNAP-CAR T cells (E:T=5:1) in a 96 well V-bottom plate with 1.0 μg/mL of the indicated BG-conjugated antibody. Plates underwent a quick-spin to collect cells at the bottom of the wells and were then incubated at 37°C for 24 hours. To identify lysed cells, co-incubated cells were stained with Ghost Dye Red Viability Dye (Tonbo Biosciences) and analyzed by flow cytometry. Target cells were identified by Cell Trace Yellow and lysed target cells were identified by positive Ghost Dye staining. Percent specific cytotoxicity was calculated by the equation: 100*(% experimental lysis – % target-only lysis) / (100 – % target-only lysis).

### Pre-labeling co-incubation assays

Pre-labeling co-incubation activation assays, were carried out as above, except prior to co-incubation for pre-labeled SNAP effector cell assays, SNAP-CAR or SNAP-synNotch Jurkat cells were first labeled with 1.0 μg/mL of the indicated BG-modified antibody in complete media for 30 minutes at 37°C and then washed three times in complete media, and for pre-labeled target cell assays, target cells were labeled with 5.0 μg/mL of the indicated antibody for 30 minutes at 4°C and washed two times with flow buffer. No additional antibody was added to these co-incubations.

### Mathematical model

The model for ternary body formation considered the following 8 binding reactions between the tumor cells, T cells, and antibody with six different species: T cell (*Tc*), antibody (*Ab*), tumor cell (*Tu*), T cell bound to antibody (*Tc.Ab*), tumor cell bound to antibody (*Ab.Tu*), and a ternary body complex of a T cell bound to antibody and tumor cell (*Tc.Ab.Tu*) and where rates *k*_*fi*_ (*i*=1..4) represent the forward kinetic rate constants, and rates *k_ri_* represent the reverse kinetic rate constants:

Reactions 1 and 2:

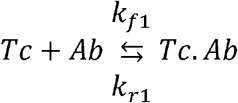

Reactions 3 and 4:

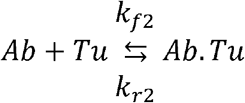

Reactions 5 and 6:

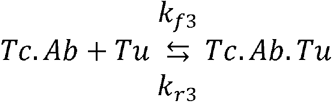

Reactions 7 and 8:

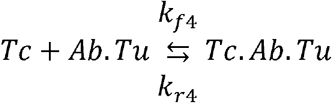

From reactions 1-8, we derived a system of equations to describe the accumulation of each of the six species in the model. In Equations 1-8 below, we list the forward and backward components of the eight reactions expressing the change in concentration of each species:

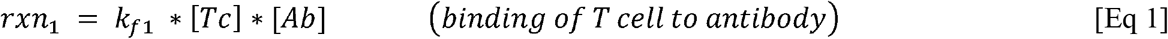

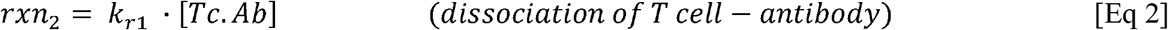

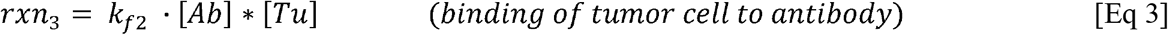

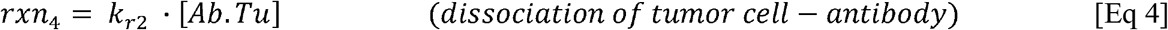

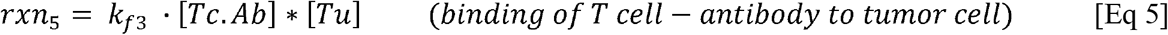

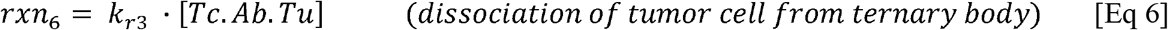

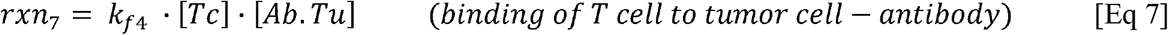

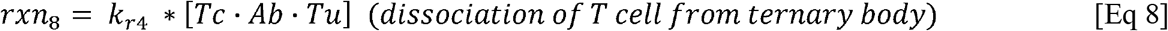

The equations 9–14 below were used to compute the change in concentration of each species.

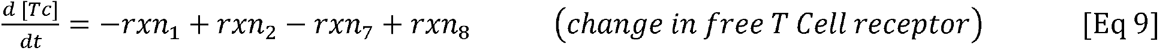

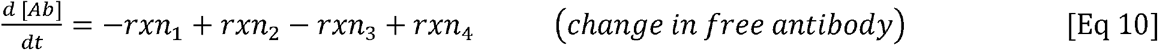

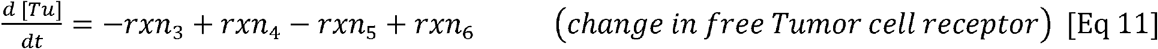

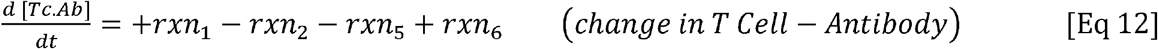

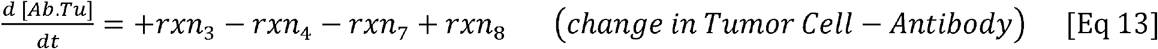

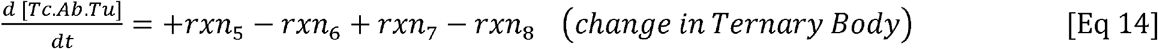

The ODE model was created under the assumption was made that the system components were well-mixed. Variables used in the ODEs were taken from the experimental design and literature values of kinetic binding and dissociation rates as summarized in **Supplementary Table S2**^34–37^. The ODE model was written in Python and solved using SciPy. To examine the concentration of each species with time, the system of ODEs was solved using the initial conditions and experimental setup values through a kinetic simulation (**Supplementary Fig. S5**). To generate equilibrium simulations (**Supplementary Fig. S6**), kinetic simulations were run for variety of antibody concentrations (10^−4^ μg/mL - 10^1^ μg/mL) and total ternary body formation from the equilibrium state of each kinetic simulation was plotted. To fit the model, we calculated the sum of squared error (SSE) between the experimental data and the simulation results. For the experimental data we used the TagBFP MFI for synNotch (**Fig. 2d**) and CD25 MFI for the read-out of SNAP-CAR activation (**Fig. 3d**). As the experimental data was only collected at specific points of antibody concentration, only the matching points in the simulations were used. Using SciPy, we minimized SSE to optimize the kinetic parameters and better fit the model to the experimental data for each antibody pair (**Fig. 6b**). During parameter estimation, the literature values were used as initial estimates and bounded within one order of magnitude. Parameter scans of *k*_*f1*_, *K*_*d*2_, and the number of tumor antigens were conducted as above for equilibrium simulations using 900 simulations over the bounds for each parameter. Ternary body formation was normalized to the maximal concentration across all simulations.

### Statistical methods

The number of replicates, mean value, and error are described in the respective figure legends and/or methods. Error bars are shown for all data points with replicates as a measure of variation within a group.

## Data availability

All data generated and analyzed during this study are included in this published article and its Supplementary Information or are available from the corresponding author upon reasonable request.

## Acknowledgments

This work was supported by the NIH grant R35 CA210039 (O.J.F.); NIH grant R21 AI130815 (A.D.); DARPA award W911NF-17-1-0135 (N.M-Z.); and by the Michael G. Wells Prize (J.L.). This work benefitted from using the SPECIAL BD LSRFORTESSA funded by NIH 1S10OD011925-01.

## Author contributions

J.L. and A.D. designed the research. A.A.B. and N.M-Z. created the mathematical model, and A.A.B. performed the model data fitting and parameter scans. J.L., Y.T., and M.K. carried out the experiments and analyzed the data. J.L., N.M-Z., A.D., and O.J.F. supervised the work. J.L., A.B., Y.T., M.K., N.M-Z., A.D., and O.J.F. interpreted the data and wrote the manuscript.

## Materials & correspondence

Correspondence and requests for materials should be addressed to J.L.

## Supplementary Information

Figure S1 | mSA2 biotin-binding synNotch receptor is incapable of cell-bound antibody activation.

Figure S2 | SDS-PAGE quantification of antibody-conjugated benzyl guanines by SNAP conjugation reaction.

Figure S3 | Staining of target cell lines by benzyl guanine modified antibodies.

Figure S4 | Effector to target effect on SNAP-synNotch receptor activity.

Figure S5 | Representative kinetic simulation. Figure S6 | Representative equilibrium simulation.

Table S1 | Quantification of the number of BG molecules conjugated per antibody.

Table S2 | Model parameters used in simulations.

Table S3 | Model simulation error.

Table S4 | DNA sequences of lentiviral receptor expression and response inserts.

