## Supplementary Information for "Post-translational covalent assembly of CAR and synNotch receptors for programmable antigen targeting"

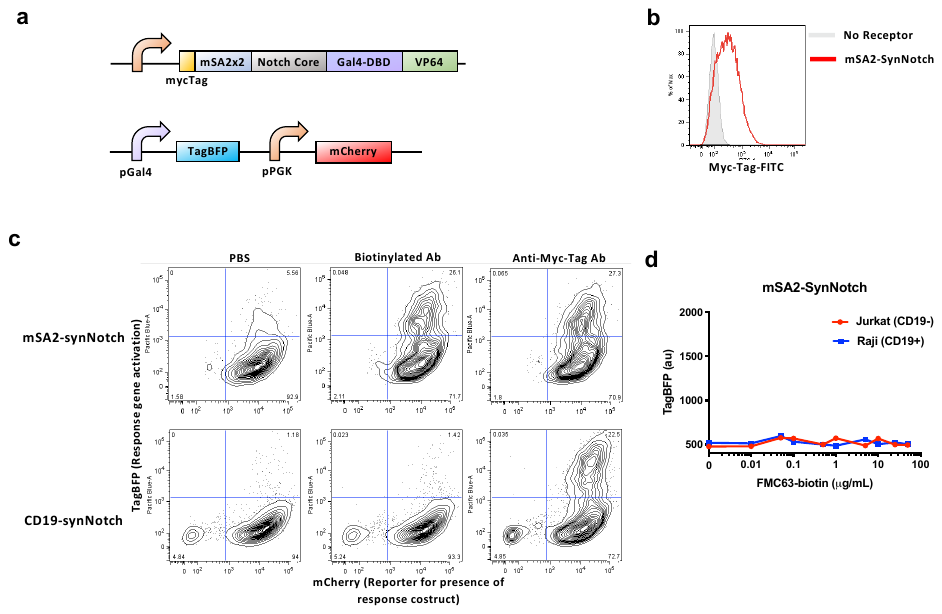

**Figure S1 | mSA2 biotin-binding synNotch receptor is activated by plate-bound biotinylated antibody but not antibody bound to the surface of target cells. a,** Design of SNAP-synNotch receptor expression and response lentiviral vectors. **b,** Flow cytometry analysis of the surface expression of the mSA2-synNotch receptor on transduced vs. MOCK (un)transduced Jurkat cells assessed by staining with the anti-Myc-Tag antibody. **c,** Flow cytometry analysis of the activation of mSA2-synNotch cells incubated on plates coated with PBS, biotinylated antibody, or anti-Myc-Tag antibody, for 48 hours for TagBFP output gene expression of response construct positive (mCherry+) cells. **d,** Flow cytometry analysis of the activation of mSA2-synNotch cells co-incubated with the indicated target cell lines and antibody concentrations for 48 hours for TagBFP output gene expression of response construct positive (mCherry+) cells reported as mean fluorescence intensity (MFI).

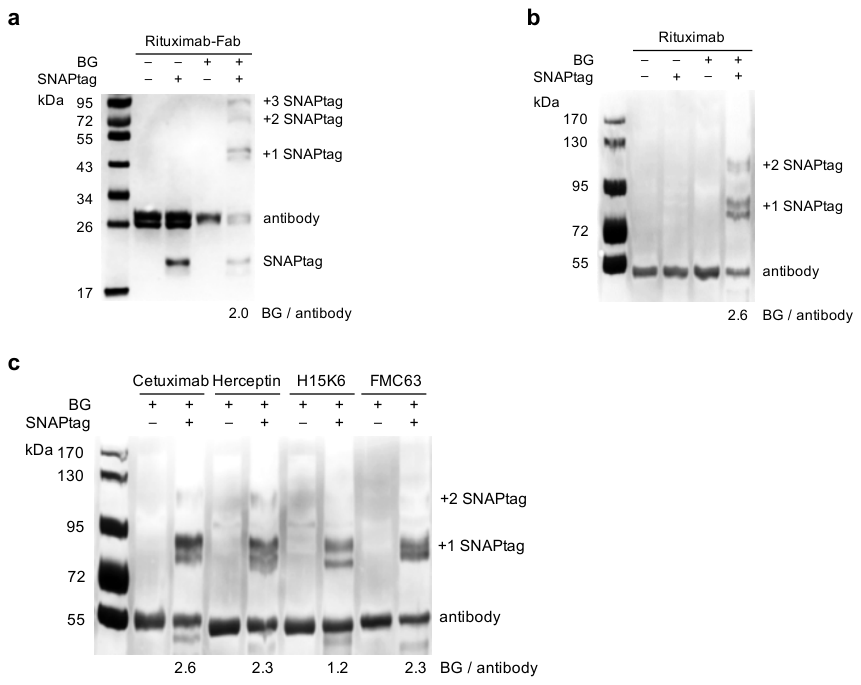

**Figure S2 | SDS-PAGE quantification of antibody-conjugated benzyl guanines by SNAP conjugation reaction.** Quantification of antibody benzylguanine (BG) labeling efficiency. SNAPtag conjugation to the light chain, **a**, or the whole antibody, **b**, was only achieved in the presence of both the BG-labeled antibody and the SNAPtag protein. A panel of BG-labeled antibodies, **c**, were then assessed in the same manner. Average number of BGs per antibody was calculated based on relative band intensities.

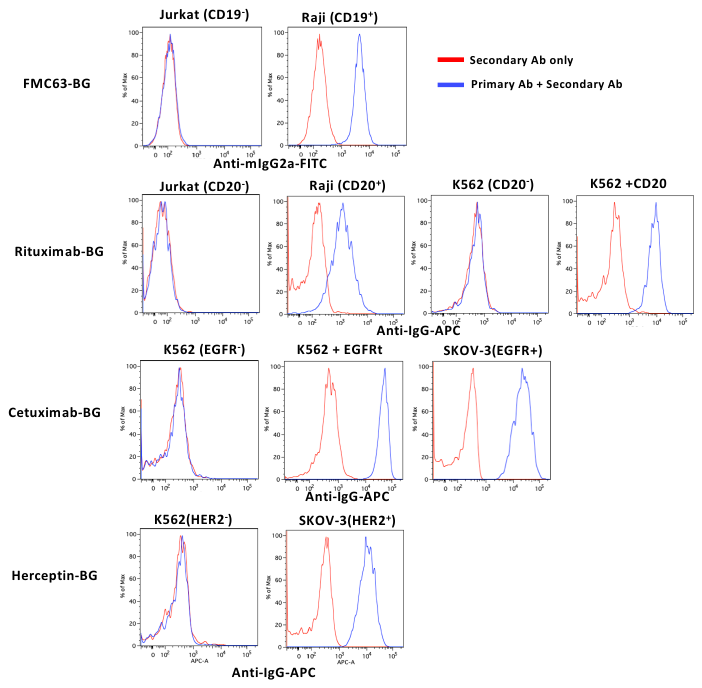

**Figure S3 | Staining of target cell lines by BG-conjugated antibodies.** Target cell lines were stained with 1.6 μg/mL of the indicated BG-conjugated antibodies (1.0 μg/mL was used for FMC63-BG) followed by staining with an anti-IgG secondary antibody (anti-mIgG2a-FITC for FMC63-BG and anti-IgG(Fab2)-APC for all other antibodies). Cells were then washed and analyzed by flow cytometry.

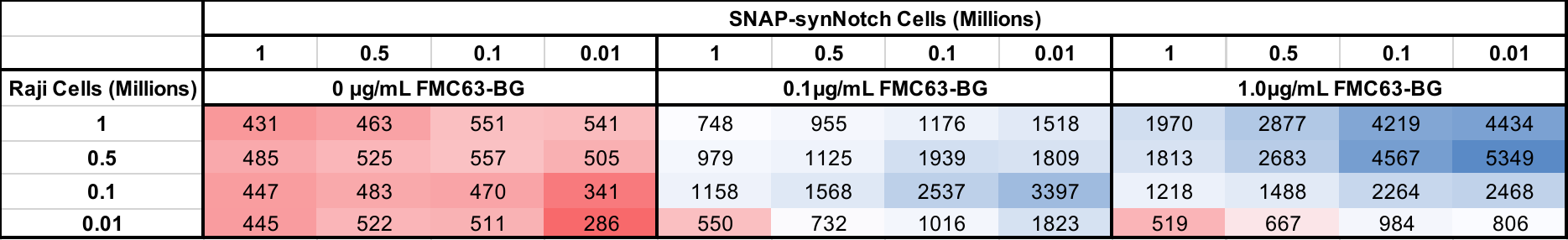

**Figure S4 | Effector to target effect on SNAP-synNotch receptor activity.** Flow cytometry analysis of the activation of SNAP-synNotch cells co-incubated with the indicated target cell lines and FMC63-BG antibody at the indicated cell numbers and antibody concentrations for 48 hours for TagBFP output gene expression reported as mean fluorescence intensity (MFI)

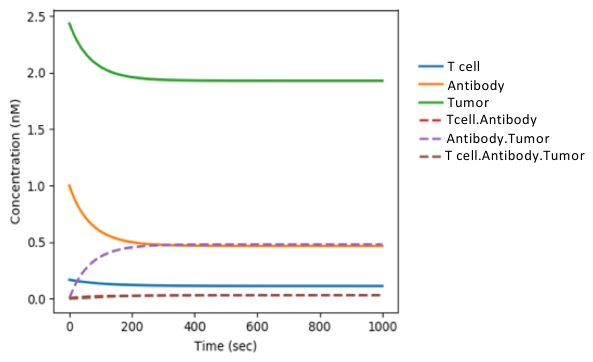

**Figure S5 | Representative kinetic simulation.** A kinetic simulation was performed using the experimental and literature values for the Cetuximab antibody and the EGFR antigen. All solid lines represent the species that are provided to the model while those in dotted lines represent complex species.

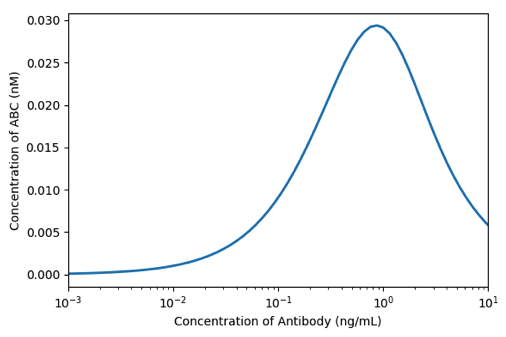
**Figure S6 | Representative equilibrium simulation.** An equilibrium simulation was performed using the experimental and literature values for the Cetuximab antibody and the EGFR antigen. “ABC” denotes the ternary body formation.

**Table S1 | Quantification of the number of BG molecules conjugated per antibody.** Data summary from *Fig. S2.*

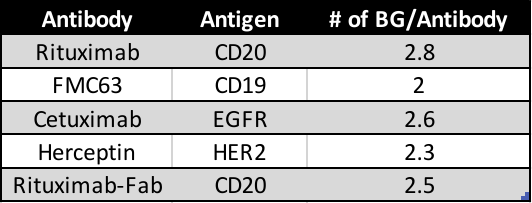

**Table S2 | Model parameters used in simulations.** Indicated parameters and their starting values as derived from the experimental set-up or from the literature.

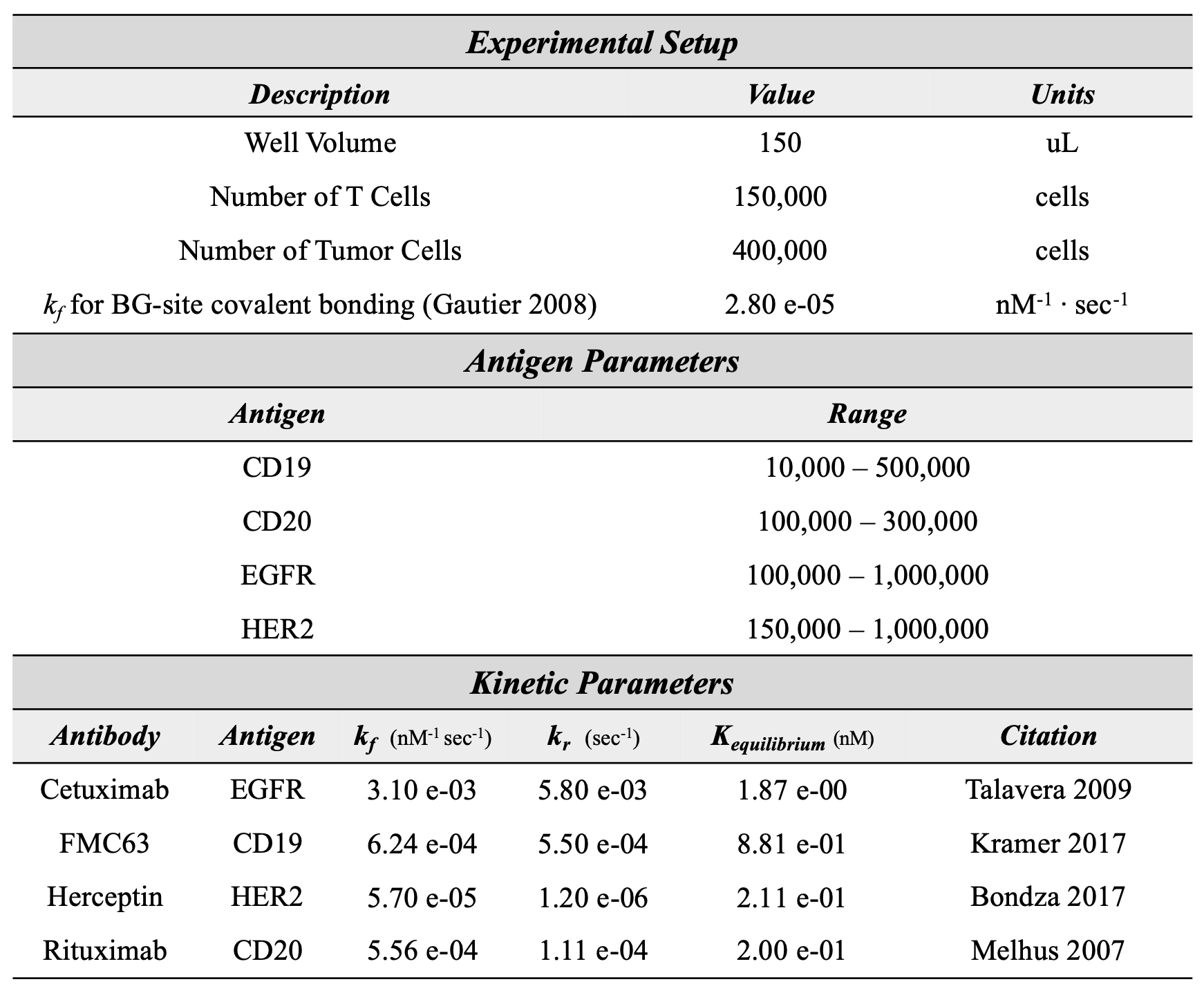

**Table S3 | Model simulation error.** Summary of sum of squared error (SSE) values calculated between the experimental results and simulations for simulations using parameters from the literature or parameters derived from model fitting.

| ***Model Error*** | | |
| --- | --- | --- |
| ***Simulation*** | ***Literature Simulation*** | ***Fitted Simulation*** |
| CAR with Cetuximab | 0.38 | 0.38 |
| CAR with FMC63 | 0.61 | 0.01 |
| CAR with Herceptin | 0.08 | 0.02 |
| CAR with Rituximab | 2.36 | 0.21 |
| synNotch with Cetuximab | 0.73 | 0.02 |
| synNotch with FMC63 | 0.20 | 0.03 |
| synNotch with Herceptin | 1.39 | 0.02 |
| synNotch with Rituximab | 2.51 | 0.02 |

**Table S4 | DNA sequences of lentiviral receptor expression and response inserts.**

**SNAP-41BBζ-T2A-TagBFP (entire coding region inserted into pHR-PGK vector)**

**GGAGCAAGGCAGGTGGACAGTGGATCatggagacagacacactcctgctatgggtgctgctgctctgggttccaggttccacaggtATGGACAAAGATTGCGAGATGAAGAGAACCACCCTGGATAGCCCTCTCGGCAAGCTCGAACTTTCTGGTTGTGAACAGGGTTTGCACAGGATCATCTTCCTGGGAAAGGGAACCTCAGCCGCAGATGCGGTTGAAGTGCCAGCTCCGGCTGCAGTGCTTGGTGGACCCGAGCCTCTTATGCAAGCAACGGCATGGCTTAATGCTTATTTTCACCAGCCTGAGGCCATTGAAGAGTTTCCAGTTCCTGCATTGCATCACCCCGTTTTTCAGCAGGAATCCTTCACTAGACAAGTGCTTTGGAAGCTCTTGAAAGTGGTTAAATTTGGGGAAGTCATCTCATACAGCCACCTTGCTGCCCTTGCAGGCAATCCTGCGGCCACGGCTGCAGTGAAAACTGCACTTAGCGGAAATCCAGTCCCCATCTTGATACCGTGTCACAGGGTAGTACAGGGCGACCTGGACGTCGGCGGTTACGAGGGCGGTTTGGCCGTTAAGGAATGGTTGCTGGCGCATGAGGGTCACCGGCTGGGAAAACCAGGTCTTGGTGGAGGAAGTGGAGGATCTaccactactccggcaccgcgccccccaactcctgcaccgacgatagcttcacaaccgctttcattgcggcccgaagcatgtcggccagccgccggaggcgctgtgcatacaagagggctggattttgcatgtgatatatatatttgggcgccccttgctggcacttgcggcgttcttcttcttagcctcgttattacgctctactgtAAGCGAGGTAGGAAAAAATTGCTGTATATCTTTAAACAGCCTTTTATGAGACCCGTGCAAACGACTCAAGAGGAAGACGGGTGTAGCTGTAGATTTCCTGAAGAGGAAGAGGGGGGGTGCGAACTGcgggtgaagttcagcagaagcgccgacgcccctgcctaccagcagggccagaatcagctgtacaacgagctgaacctgggcagaagggaagagtacgacgtcctggataagcggagaggccgggaccctgagatgggcggcaagcctcggcggaagaacccccaggaaggcctgtataacgaactgcAgaaagacaagatggccgaggcctacagcgagatcggcatgaagggcgagcggaggcggggcaagggccacgacggcctgtatcagggcctgtccaccgccaccaaggatacctacgacgccctgcacatgcaggccctgcccccaaggCTCGAGGGCGGCGGAGAGGGCAGAGGAAGTCTTCTAACATGCGGTGACGTGGAGGAGAATCCCGGCCCTCGCatgagcgagctgattaaggagaacatgcacatgaagctgtacatggagggcaccgtggacaaccatcacttcaagtgcacatccgagggcgaaggcaagccctacgagggcacccagaccatgagaatcaaggtggtcgagggcggccctctccccttcgccttcgacatcctggctactagcttcctctacggcagcaagaccttcatcaaccacacccagggcatccccgacttcttcaagcagtccttccctgagggcttcacatgggagagagtcaccacatacgaagacgggggcgtgctgaccgctacccaggacaccagcctccaggacggctgcctcatctacaacgtcaagatcagaggggtgaacttcacatccaacggccctgtgatgcagaagaaaacactcggctgggaggccttcaccgagacgctgtaccccgctgacggcggcctggaaggcagaaacgacatggccctgaagctcgtgggcgggagccatctgatcgcaaacatcaagaccacatatagatccaagaaacccgctaagaacctcaagatgcctggcgtctactatgtggactacagactggaaagaatcaaggaggccaacaacgaaacatacgtcgagcagcacgaggtggcagtggccagatactgcgacctccctagcaaactggggcacaagcttaattaaGATCCTTGACTTGCGGCCGCAACTCCCAC**

**Legend:**

**5’ region of pHR-PGK acceptor vector**

**Leader sequence and Kozak sequence**

**SNAPtag**

**CD8α-hinge,TM**

**4-1BBcyto**

**CD3ζcyto**

**T2A**

**TagBFP**

**3’ region of pHR-PGK acceptor vector**

**pHR-PGK-SNAP-synNotch-Gal4-VP64 (insert for pHR–PGK-[]-synNotch-Gal4-VP64 backbone)**

**gagcaaaaacttatctctgaagaggacctcATGGACAAAGATTGCGAGATGAAGAGAACCACCCTGGATAGCCCTCTCGGCAAGCTCGAACTTTCTGGTTGTGAACAGGGTTTGCACAGGATCATCTTCCTGGGAAAGGGAACCTCAGCCGCAGATGCGGTTGAAGTGCCAGCTCCGGCTGCAGTGCTTGGTGGACCCGAGCCTCTTATGCAAGCAACGGCATGGCTTAATGCTTATTTTCACCAGCCTGAGGCCATTGAAGAGTTTCCAGTTCCTGCATTGCATCACCCCGTTTTTCAGCAGGAATCCTTCACTAGACAAGTGCTTTGGAAGCTCTTGAAAGTGGTTAAATTTGGGGAAGTCATCTCATACAGCCACCTTGCTGCCCTTGCAGGCAATCCTGCGGCCACGGCTGCAGTGAAAACTGCACTTAGCGGAAATCCAGTCCCCATCTTGATACCGTGTCACAGGGTAGTACAGGGCGACCTGGACGTCGGCGGTTACGAGGGCGGTTTGGCCGTTAAGGAATGGTTGCTGGCGCATGAGGGTCACCGGCTGGGAAAACCAGGTCTTGGTGGAGGAAGTGGAGGATCTatcctggactacagcttcacaggtggcgct**

**Legend:**

**5’ region of pHR_PGK_antiCD19_synNotch_Gal4VP64 acceptor vector**

**SNAPtag**

**3’ region of pHR_PGK_antiCD19_synNotch_Gal4VP64 acceptor vector**

**pHR-PGK-mSA2x2-synNotch-Gal4-VP64 (insert for pHR_PGK_[]_synNotch_ Gal4VP64 vector backbone)**

**gagcaaaaacttatctctgaagaggacctcGGCGCGGAGGCTGGAATCACTGGCACTTGGTATAATCAGCATGGCTCTACCTTTACTGTAACAGCAGGGGCTGATGGCAACCTTACCGGCCAATACGAGAATCGGGCGCAGGGTACGGGATGTCAAAATTCTCCCTATACACTGACCGGACGGTATAACGGTACCAAACTCGAATGGAGGGTAGAATGGAACAACTCTACAGAAAACTGCCATAGCAGAACCGAATGGAGAGGCCAGTATCAAGGAGGTGCAGAAGCCCGAATTAACACCCAATGGAACTTGACTTACGAAGGCGGATCTGGACCAGCGACGGAACAGGGGCAGGATACTTTCACCAAAGTAAAGCCAAGTGCTGCGTCCGGTAGTGAGGCCGCTGCTAAAGAAGCAGCGGCCAAGGAAGCAGCTGCTAAGGGTGCAGAGGCTGGTATAACGGGGACTTGGTACAACCAGCACGGCTCCACCTTTACTGTGACTGCTGGGGCAGATGGGAACTTGACAGGACAATATGAGAACCGAGCACAAGGGACGGGCTGCCAAAATAGTCCATATACACTCACGGGGCGATACAATGGCACTAAGCTGGAATGGAGGGTTGAGTGGAACAATTCAACGGAAAACTGTCATTCCCGCACTGAATGGCGGGGGCAGTACCAGGGGGGGGCGGAGGCGAGAATCAACACACAATGGAACTTGACATACGAGGGGGGAAGTGGGCCTGCCACCGAACAAGGACAGGACACTTTTACTAAAGTGAAGCCCTCAGCTGCGTCAGGGAGTatcctggactacagcttcacaggtggcgct**

**Legend:**

**5’ region of pHR_PGK_antiCD19_synNotch_Gal4VP64 acceptor vector**

**mSA2x2**

**3’ region of pHR_PGK_antiCD19_synNotch_Gal4VP64 acceptor vector**

**pHR-pGal-IL7-PGK-mCherry (insert for pHR_Gal4UAS_[]_PGK_mCherry vector backbone)**

**CCGATCCAGCCTCTCGACATTCGTTGGATCatgttccacgtaagtttcagatatatctttggacttccgccgctcatattggtattgttgccagtggcatctagtgactgtgacatagaaggaaaggatggtaaacagtatgaaagcgtacttatggtatccattgaccagcttctcgatagtatgaaagagattggtagtaattgcctcaataacgagttcaatttctttaaacgacacatttgtgatgcgaataaagagggaatgtttctgtttcgcgccgcgaggaagcttaggcagttccttaaaatgaactcaactggggatttcgacctccatctgctgaaggtgagtgaaggtactactattctcctgaattgcacgggacaggtaaaggggcgaaaacctgcggccttgggtgaggcacaaccaaccaaaagcctcgaagaaaacaagtccctcaaagaacagaagaagctcaacgatctgtgctttctgaaaagactcttgcaggagatcaaaacttgttggaataagattttgatgggcactaaggagcatTAAGATCCTTGACTTGCGGCCGCAACTCCC**

**Legend:**

**5’ region of pHR_Gal4UAS_[]_PGK_mCherry acceptor vector**

**IL-7**

**3’ region of pHR_PGK_antiCD19_synNotch_Gal4VP64 acceptor vector**
